# Draft Genome Sequence of pyschrotolerant *Clostridium* sp. M14 Isolated from Spoiled Uncooked Venison

**DOI:** 10.1101/2020.03.24.006783

**Authors:** Nikola Palevich, Faith P. Palevich, Paul H. Maclean, Ruy Jauregui, Eric Altermann, John Mills, Gale Brightwell

## Abstract

The non-proteolytic, non-toxigenic and pyschrotolerant *Clostridium* sp. M14 was isolated from vacuum-packaged refrigerated spoiled venison. This report describes the generation and annotation of the 3.9 Mb draft genome sequence of *Clostridium* sp. M14.

## ANNOUNCEMENT

The pyschrotolerant *Clostridium* sp. M14 is a Gram-positive, rod-shaped, spore-forming and slow-growing obligate anaerobe. M14 is able to grow at temperatures below 4°C, is catalase□ and oxidase□negative and metronidazole□sensitive, and originally isolated from vacuum-packaged refrigerated spoiled venison (1). Numerous bacterial species belonging to the *Clostridium* genera have been recognized as causative agents of blown pack spoilage (BPS) in vacuum packed meat products (2), and M14 was selected for genome sequencing to examine its role in BPS.

Previous studies identified a number of isolates with some similarity to that of non-proteolytic *C. botulinum* type B using classical bio-chemical differentiation, 16*S* rRNA gene Restriction Fragment Length Polymorphism (RFLP) pattern analyses, 16S rRNA gene sequencing and PCR amplification of the ITS regions (1, 3). However, none of these isolates were deemed toxigenic. Broda *et al*., concluded that although the growth of such microorganisms in vacuum□packed chilled meat leads to product spoilage, it does not prejudice product safety. Here, we present a draft genome sequence of a representative meat spoilage associated non-proteolytic and non-toxigenic pyschrotolerant *Clostridium* sp. M14, that falls within meat associated pyschrotolerant *Clostridium* ARDRA Group 7 (4).

Strain M14 was isolated from a fully blown pack of vacuum packaged venison nearly 20 years ago and cultured anaerobically at 10°C in pre-reduced Peptone, Yeast Extract, Glucose, Starch broth (PYGS) as previously described (5). Genomic DNA was extracted using a modified phenol-chloroform procedure (5) and mechanically sheared using a Nebulizer instrument (Invitrogen) to select fragments of approximately 550 bp. A DNA library was prepared using the Illumina TruSeq™ Nano method and sequenced on the Illumina MiSeq platform with the 2× 250 bp paired-end (PE) reagent kit v2 producing a total of 3,125,724 PE raw reads. The quality of the raw reads was checked in FastQC v0.11.5 (https://www.bioinformatics.babraham.ac.uk/projects/fastqc/), the reads were trimmed with Trimmomatic v0.39 (http://www.usadellab.org/cms/?page=trimmomatic) and assembled using the A5-miseq pipeline v20169825 with standard parameters (6). The *de novo* assembly of M14 produced 36 scaffolds with 184× coverage and an N50 value of 757,921 bp, with the largest scaffold length being 1,669,648 bp in size. The draft genome sequence is composed of 3,986,879 bp, with a %G+C content of 27.1%. A total of 3,717 putative protein-coding genes (PCGs) were predicted along with 81 tRNA, 19 rRNA, 170 ncRNA elements using GAMOLA2 (7). In addition, Diamond v0.9.21.122 (8) and InterProScan v5.36-75.0 (9) were used to search the NCBI “nr” database with the resulting protein set imported into BLAST2GO as implemented in the OmicsBox software package v1.1.164 (10) where gene ontology terms and final annotations were assigned to each protein. All bioinformatics analyses were performed using default settings and parameters.

Carbohydrate-Active enZYme (11) profiling was analyzed using dbCAN2 (12) and revealed that the M14 genome is predicted to encode 44 glycoside hydrolases (GHs), 27 glycosyl transferases (GTs), 10 carbohydrate esterases (CEs) and 19 carbohydrate-binding protein module (CBM) families. Overall, approximately 2.7% of the M14 genome (100 CDSs) is predicted to encode either secreted or intracellular proteins dedicated to carbohydrate and even polysaccharide degradation. A comparison of the M14 with the recently characterized *C. estertheticum* subsp. *laramiense* type strain DSM 14864^T^ (ATCC 51254) (5), revealed clear differences in their enzymatic profiles. While both appear to be equipped to utilize many oligo- and monosaccharides as substrates for growth and encode a large repertoire of enzymes predicted to metabolize complex insoluble polysaccharides, M14 lacked genes encoding polysaccharide lyases (PLs).

The non-toxigenic status of M14 was confirmed by performing a bioinformatics search of the whole genome sequence (WGS) database (visited March 2020) at the National Center for Biotechnology Information (NCBI), using the predicted protein translation products of the known deadly neurotoxins of *C. botulinum* (BoNTs) protein sequences. None of the putative neurotoxin genes within the Group II *C. botulinum* BoNT operon that consists of *ha* (haemagglutinin) types (*ha70, ha17* and *ha34*) and accessory proteins *ntnh* (nontoxic-nonhaemagglutinin) and *botR* (neurotoxin regulator protein) upstream of the boNT gene, were identified in M14. In addition, a whole-genome alignment to the publicly available Group II *C. botulinum* genomes revealed a low sequence homology with approximately <20% similarity to M14.

The genome sequence of the pyschrotolerant *Clostridium* sp. M14 reported here is a valuable resource for future studies investigating the bacterial genetic mechanisms associated with BPS. In order to improve the phylogenetic resolution of the *Clostridium* genera and improve our limited knowledge of meat spoilage caused by non-toxigenic *Clostridia* species.

### Data availability

The genome sequence and associated data for *Clostridium* sp. M14 were deposited under GenBank accession number JAAMNF000000000, BioProject accession number PRJNA574489 and in the Sequence Read Archive (SRA) under accession number SRR11113219.

